# CK2 phosphorylation of human papillomavirus 16 E2 on serine 23 promotes interaction with TopBP1 and is critical for E2 plasmid retention function

**DOI:** 10.1101/2021.02.17.431757

**Authors:** Apurva T. Prabhakar, Claire D. James, Dipon Das, Raymonde Otoa, Matthew Day, John Burgner, Christian T. Fontan, Xu Wang, Andreas Wieland, Mary M. Donaldson, Molly L. Bristol, Renfeng Li, Anthony W. Oliver, Laurence H. Pearl, Brian O. Smith, Iain M. Morgan

**Affiliations:** Virginia Commonwealth University (VCU), Philips Institute for Oral Health Research, School of Dentistry, Richmond, VA, 23298; Cancer Research UK DNA Repair Enzymes Group, Genome Damage and Stability Centre, School of Life Sciences, University of Sussex, Brighton, United Kingdom; Emory Vaccine Center and Department of Microbiology & Immunology, Emory University School of Medicine, Atlanta, GA, 30322; School of Veterinary Medicine, University of Glasgow, Garscube Estate, Bearsden, UK; VCU Massey Cancer Center, Richmond, VA, 23298; Institute of Molecular, Cell & Systems Biology, University of Glasgow, Glasgow, G12 8QQ, UK

**Keywords:** human papillomavirus, E2, TopBP1, BRD4, segregation, retention, cervical cancer, head and neck cancer, life cycle, CK2, phosphorylation, assay

## Abstract

During the human papillomavirus 16 (HPV16) life cycle, the E2 protein interacts with host factors to regulate viral transcription, replication and genome segregation/retention. Our understanding of host partner proteins and their roles in E2 functions remains incomplete. Here, we demonstrate that CK2 phosphorylation of E2 on serine 23 promotes interaction with TopBP1 *in vitro* and *in vivo*, and that E2 is phosphorylated on this residue during the HPV16 life cycle. We investigated the consequences of mutating serine 23 on E2 functions. E2-S23A activates and represses transcription identically to E2-WT (wild-type), and E2-S23A is as efficient as E2-WT in transient replication assays. However, E2-S23A has compromised interaction with mitotic chromatin when compared with E2-WT. In E2-WT cells, both E2 and TopBP1 levels increase during mitosis when compared with vector control cells. In E2-S23A cells, neither E2 nor TopBP1 levels increase during mitosis. We next tested whether this difference in E2-S23A levels during mitosis disrupts E2 plasmid retention function. We developed a novel plasmid retention assay and demonstrate that E2-S23A is deficient in plasmid retention when compared with E2-WT. siRNA targeted knockdown of TopBP1 abrogates E2-WT plasmid retention function. Introduction of the S23A mutation into the HPV16 genome resulted in delayed immortalization of human foreskin keratinocytes (HFK) and higher episomal viral genome copy number in resulting established HFK. Overall, our results demonstrate that CK2 phosphorylation of E2 on serine 23 promotes interaction with TopBP1, which is critical for E2 plasmid retention function and in HPV16 immortalization of keratinocytes.

**Importance:** Human papillomaviruses are causative agents in around 5% of all cancers, with no specific anti-viral therapeutics available for treating infections or resultant cancers. In this report, we demonstrate that phosphorylation of HPV16 E2 by CK2 promotes formation of a complex formation with the cellular protein TopBP1 *in vitro* and *in vivo*. This complex results in stabilization of E2 during mitosis and mediates plasmid retention by E2. This function promotes the partitioning of viral genomes into the nuclei of daughter cells following mitosis. We demonstrate that CK2 phosphorylates E2 on serine 23 *in vivo*, and that CK2 inhibitors disrupt the E2-TopBP1 complex. Mutation of E2 serine 23 to alanine disrupts the HPV16 life cycle, demonstrating a critical function for this residue. Together, our results suggest that CK2 inhibitors may disrupt the E2-TopBP1 dependent HPV16 life cycle and potentially kill HPV16 positive cancers, which lays a molecular foundation to develop novel therapeutic approaches for combating HPV16 disease.

## Introduction

Human papillomaviruses (HPVs) infection leads to around 5% of all human cancers, with HPV16 infection being responsible for 50% of cervical cancers and 80-90% of HPV positive oropharyngeal cancers (1). This latter disease has increased dramatically in the last generation and represents an ongoing public health crisis with no specific anti-viral therapeutics available for combating the disease (2-6). Identification of such therapeutics are a priority, and our lab focuses on enhancing the molecular understanding of the HPV16 life cycle in order to identify potential anti-viral targets.

HPVs infect basal epithelial cells and following cell division the viral DNA locates to the cell nucleus (7). The viral genome then replicates to 20-50 copies per cell and the infected cell begins to proliferate, promoted by the expression of E6 and E7 that target p53 and pRb (among other proteins), respectively (8, 9). During proliferation, the viral genome copy number is maintained at around 20-50 copies per cell and in the upper layers of the epithelium there is a replication amplification stage where the viral genome copy number increases. The viral structural proteins L1 and L2 are then expressed and encapsulate the viral DNA to form viral particles that egress from the upper layers of the epithelium (10-12).

Throughout the viral life cycle there are two viral proteins that mediate replication of the viral genome, E1 and E2 (13-15). E2 is a DNA binding factor whose carboxyl terminal domain forms homo-dimers that bind to three 12bp palindromic DNA sequences surrounding the viral origin of replication in the long control region (LCR), adjacent to the transcriptional start site (16, 17). There is a fourth E2 target site in the LCR further upstream from the viral origin of replication. Following binding to its target sequences, E2 recruits the viral helicase E1 to the origin of replication via a protein-protein interaction (14-16). At the A/T rich origin E1 forms a di-hexameric helicase complex that interacts with host DNA polymerases to initiate viral DNA replication (18-21). Furthermore, E2 has additional roles during the viral life cycle. E2 can regulate transcription from the viral genome and can either activate or repress transcription depending upon the E2 concentration (22). E2 also regulates transcription from the host genome, and this regulation is directly relevant to the viral life cycle (23, 24). The fourth function for E2 during the viral life cycle is to mediate viral genome segregation (25). During cell division, the 8kbp episomal viral genome could be excluded from the nuclei of resulting daughter cells. To combat this, the virus has an active mechanism that retains viral genomes in daughter nuclei via hitch-hiking onto the host chromatin during mitosis (25). E2 mediates this function by binding to the viral DNA via its carboxyl terminal DNA binding domain and simultaneously binding to host chromatin via the E2 amino terminal domain (25). For Bovine Papillomavirus 1 (BPV1) E2, the host receptor mediating interaction with mitotic chromatin is BRD4 (26-29). For high-risk HPV (HR-HPV, those that cause cancer, including HPV16), the E2 proteins do not co-localize with BRD4 on mitotic chromatin, indicating that BRD4 may not be the mitotic receptor for these E2 proteins (16, 30, 31). We identified TopBP1 as a functional interacting partner for HPV16 E2 (32-36). TopBP1 regulates the interaction of E2 with host chromatin in interphase cells, and co-localizes with TopBP1 on mitotic chromatin, indicating that TopBP1 is a candidate protein for mediating E2 interaction with mitotic chromatin (37).

TopBP1 is a multi-functional protein involved in several aspects of nucleic acid metabolism (38). It is part of the replication complex in mammalian cells, interacting with Treslin to promote the initiation of replication (39-43). TopBP1 encodes 9 BRCT (<u>BR</u>CA1 <u>c</u>arboxyl <u>t</u>erminal) domains that act as hydrophobic pockets mediating interaction with cellular proteins, including proteins that are phosphorylated following cell signaling events and are involved in replication initiation and the DNA damage response (DDR) (44-65). TopBP1 is required for the activation of the ATR kinase via interaction with ATRIP (ATR interacting protein) and TopBP1 is also a substrate for ATM. Both ATM and ATR are activated during the viral life cycle in order to promote viral genome replication, therefore TopBP1 is an essential mediator of the HPV16 life cycle (66-70). TopBP1 also has several roles during mitosis as it prevents transmission of DNA damage (including DNA double strand breaks and catenated DNA) to G1 daughter cells (45, 47, 65, 71-74).

Our previous work identified a mutant of E2 that had a compromised interaction with TopBP1, asparagine 89 and glutamic acid 90 of E2 were mutated to tyrosine and valine, respectively (36). The change in nature from polar and charged to bulkier and more hydrophobic at the substituted residues disrupted the interaction between HPV16 E2 and TopBP1 (from now on, E2 will mean HPV16 unless stated otherwise). To gain a more mechanistic understanding of the E2-TopBP1 interaction, and how it is regulated, we tested potential phosphorylation sites on E2 that mediate TopBP1 interaction, as TopBP1 binds a number of phosphorylated proteins via its BRCT domains (38). Here we demonstrate that CK2 phosphorylation of E2 serine 23 promotes the interaction between E2 and TopBP1 *in vitro* and *in vivo*. E2-S23A (an alanine substitution at position serine 23 which is defective in TopBP1 interaction) and E2-WT have similar transcription and replication functions in our transient assays. We describe a novel plasmid segregation assay and demonstrate that E2-S23A is deficient in this function when compared with E2-WT, potentially due to a reduction of E2-S23A protein levels during mitosis when compared with E2-WT. Introduction of the E2-S23A mutation into the HPV16 genome results in a delay in human foreskin keratinocyte (HFK) immortalization when compared with the wild type genome. However, cells with the E2-S23A mutant genomes did grow out and had an elevated level of episomal viral genomes when compared with the wild type genome. Our results demonstrate a critical role for CK2 phosphorylation of E2-S23 that is important for E2 plasmid retention function and immortalization of HFK; the implications of these findings will be discussed.

## Results

### E2 serine 23 is critical for TopBP1 interaction *in vivo*

Because TopBP1 binds to phosphorylated proteins, we investigated the ability of potential phosphorylation sites on E2 to mediate the interaction with TopBP1. E2 protein sequence analysis showed that serine 23 is highly conserved in alpha-type HPV (that incorporate high-risk, HR-HPV) (Figure 1A). Also, on the crystal structure model for HPV16 E2 serine 23 juxtaposes with amino acids 89 and 90, mutation of which disrupts E2-TopBP1 interaction (36, 75). To investigate the interaction between E2 and TopBP1 via serine 23, U2OS cells stably expressing E2-WT (wild-type), E2-S23A (serine mutated to alanine) and E2-S23D (serine mutated to aspartic acid) were generated, along with pcDNA empty vector plasmid control (Figure 1B). Cell extracts from Figure 1B were immunoprecipitated by a TopBP1 antibody and TopBP1 and E2 were detected using western blotting (Figure 1C); this experiment was repeated on three independent extracts and quantitated (Figure 1D). While E2-WT and E2-S23D co-precipitate with TopBP1 (lanes 4 and 5, Figure 2C), E2-S23A is significantly compromised in this interaction (lane 3 Figure 1C, and Figure 1D).

**Figure 1.**
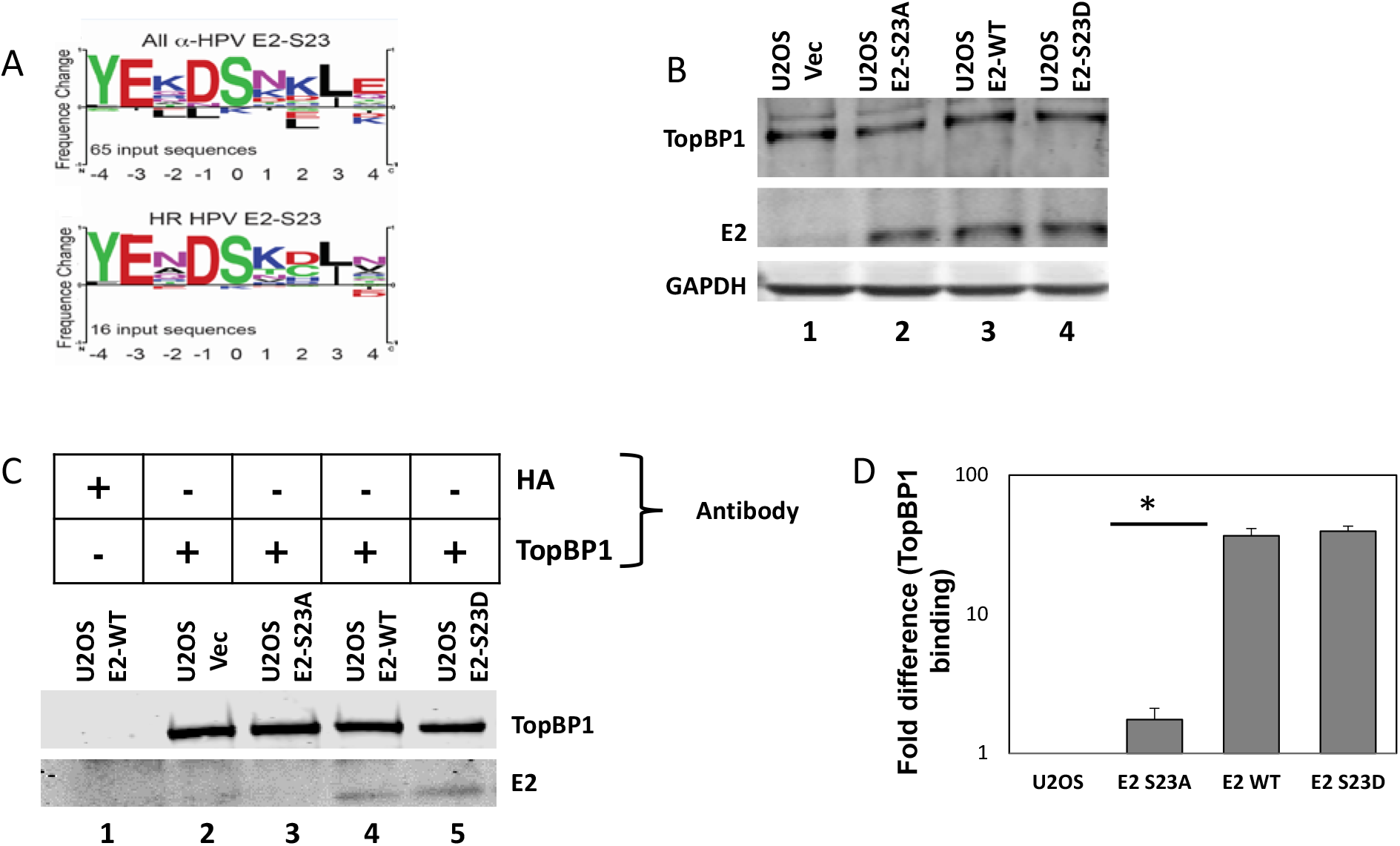
A. Motif analysis of the E2 serine 23 residue region in all α-HPV (top) and high-risk (HR, cancer causing) HPV (bottom). B. Western blots of U2OS cells expressing the indicated E2 proteins. C. Immunoprecipitation with HA (control) and TopBP1 antibody followed by western blotting for TopBP1 and E2. D. Quantitation of repeat TopBP1 co-IPs. * indicates a significant decrease in E2-S3A interaction with TopBP1 when compared with E2-WT, p-value < 0.05.

**Figure 2.**
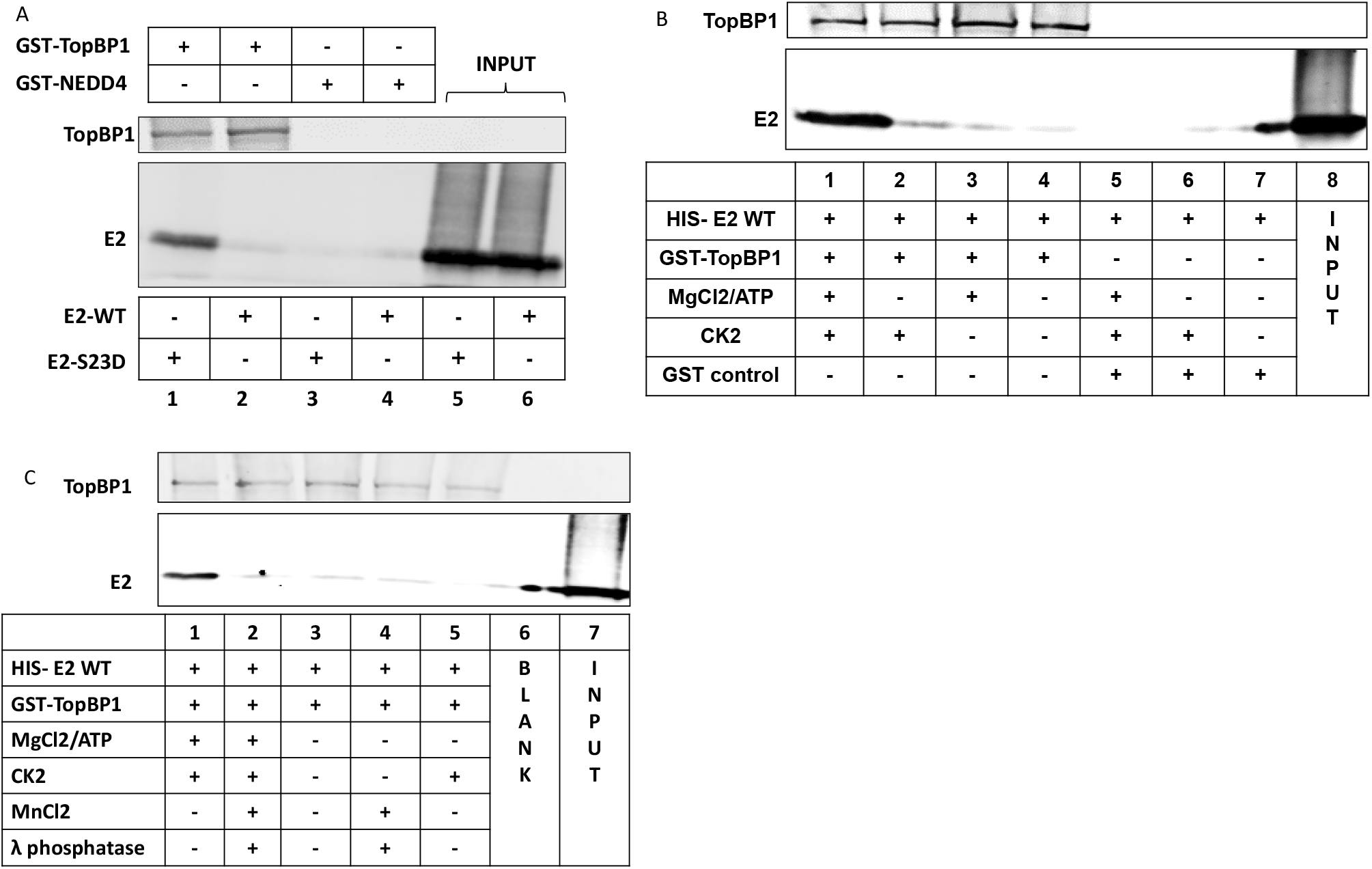
A. 0.65 pmol of GST-TopBP1 or GST-NEDD4 were incubated with 11 pmol E2 and incubated at 4°C for one hour with rotation. GST pull-downs followed by western blotting for TopBP1 (top panel) and E2 (bottom panel) were then carried out. B. The GST pull down was repeated as in A following 1 hour at 30^0^C with CK2 and controls. C. Lambda phosphatase was added to the CK2 reaction and GST pull-down assays carried out as in A. Figures S1A-C summarizes quantitation of repeat experiments. In the input lanes, only E2 was added, the TopBP1 pull down demonstrates equivalent levels of TopBP1 in each reaction containing TopBP1.

### CK2 phosphorylation of E2 promotes interaction with TopBP1 *in vitro* and *in vivo*

The negative charges at position −1 and −3 in the E2 consensus sequence around serine 23 (Figure 1A) indicate a potential CK2 target site at serine 23 (76). CK2 is also active during mitosis and could therefore be involved in mediating the plasmid retention function of E2 (77). CK2 interacts with E2, and TopBP1 interacts with CK2 phosphorylated proteins, therefore we investigated whether CK2 phosphorylates E2 serine 23 (59, 78-80). We prepared recombinant GST-TopBP1 (full-length), His-E2-WT and His-E2-S23D (amino acids 1-200 for both E2 proteins) from bacteria. These were incubated together and GST pull-down experiments carried out followed by western blotting (Figure 2A). Lanes 5 and 6 demonstrate equivalent levels of E2-S23D and E2-WT input in the GST interactions. Lane 1 demonstrates an interaction between E2-S23D and GST-TopBP1, while lane 2 demonstrates that E2-WT does not interact with TopBP1. Neither protein interacts with the GST-NEDD4 control protein. This experiment was repeated and the results quantitated (Figure S1A). To determine whether CK2 phosphorylation of E2-WT can promote interaction with TopBP1, we incubated the recombinant proteins with CK2 enzyme prior to the GST pull-down experiments (Figure 2B). Lane 1 demonstrates that the presence of an enzymatically active CK2 promotes interaction between E2-WT and TopBP1. Omission of CK2 co-factors (MgCl2/ATP) (lane 2) or CK2 enzyme (lane 3) abolished the interaction. CK2 did not promote interaction with GST-NEDD4 (present in lanes 5-7). This experiment was repeated and the results quantitated (Figure S1B). To confirm that it is CK2 phosphorylation promoting the interaction between E2-WT and TopBP1, we repeated the experiment in Figure 2B in the presence of Lambda phosphatase which eliminates the interaction between E2 and TopBP1 (Figure 2C, lane 2). This experiment was repeated and the results quantitated (Figure S1C).

To investigate whether E2 S23 is phosphorylated *in vivo* by CK2, we generated an antibody specific for phosphorylated serine 23 (pS23-Ab) using a phospho-peptide incorporating the region around S23 with the serine phosphorylated (CKILTHYENDS^P^TDLR). To investigate whether CK2 was responsible for phosphorylating serine 23 *in vivo*, we knocked down CK2 components using siRNA. Figure 3A demonstrates that expression of CK2α and α’ was down-regulated using siRNA (lanes 2 and 3, respectively), both were also targeted (lane 4). Non-specific scrambled control siRNA (Scr) was used as a control for siRNA treatment in U2OS-Vec and U2OS-E2-WT (lanes 1 and 5 respectively). When both siRNAs were combined there was a partial knock down of both proteins. There was visible toxicity in the double knockdown cells and the partial knockdown is likely due to a survival advantage of cells exhibiting a lower degree of both CK2 component knockdown. Following immunoprecipitation with pS23-Ab, there is co-immunoprecipitation (co-IP) of E2 in the Scr treated U2OS-E2-WT cells (lane 5, lower panel). In all conditions when CK2 components are targeted by specific siRNAs, there is an abrogation of detectable E2 co-IP with the pS23-Ab (lanes 2-4). To confirm the role of CK2 in phosphorylating E2 S23 to promote interaction with TopBP1, we used the CK2 inhibitor CX4945 (Figure 3B). In the presence of CX4945 (lanes 3-4) the interaction between E2 and TopBP1 is disrupted, and pS23-Ab fails to co-IP E2. To confirm that knockdown of CK2 components disrupts the E2-TopBP1 interaction we carried out TopBP1 co-IPs following CK2 α or α’ siRNA knockdown. Figure 3C (lanes 2 and 6 are blank) demonstrates that knockdown of CK2α reduces the interaction between E2 and TopBP1. There is knockdown of CK2α expression with the targeting siRNA, while Scr (scrambled) control has robust CK2α expression (compare lane 4 with 1 and 3). The HA control antibody IP does not immunoprecipitate TopBP1 or E2 (lane 5), while TopBP1 antibody pulls down both proteins (lanes 7 and 8). There is a reduced co-IP of E2 with the TopBP1 antibody when CK2α is knocked-down (compare lane 7 with lane 8). This experiment was repeated and quantitated and there is a statistically significant reduction in the interaction between TopBP1 and E2 following knockdown of CK2α expression (Figure S2A). Figure 3D demonstrates that knockdown of CK2α’ expression also reduces the E2-TopBP1 interaction. Lane 3 (taken from the same gel as lanes 1 and 2 with lanes removed) demonstrates a reduction of CK2α’ with targeting siRNA when compared with the control Scr siRNA. Immunoprecipitation with TopBP1 antibody results in a co-IP with E2 (lanes 5 and 6). As with CK2α, there is a reduction in the amount of E2 co-IPed with TopBP1 when CK2α’ is knocked-down (compare lane 5 with lane 6). These experiments were repeated and quantitated and there is a statistically significant reduction in the interaction between TopBP1 and E2 following knockdown of CK2α’ expression (Figure S2B). The E2-TopBP1 interaction is not completely eliminated because CK2α and CK2α’ likely compensate for the absence of the other.

**Figure 3.**
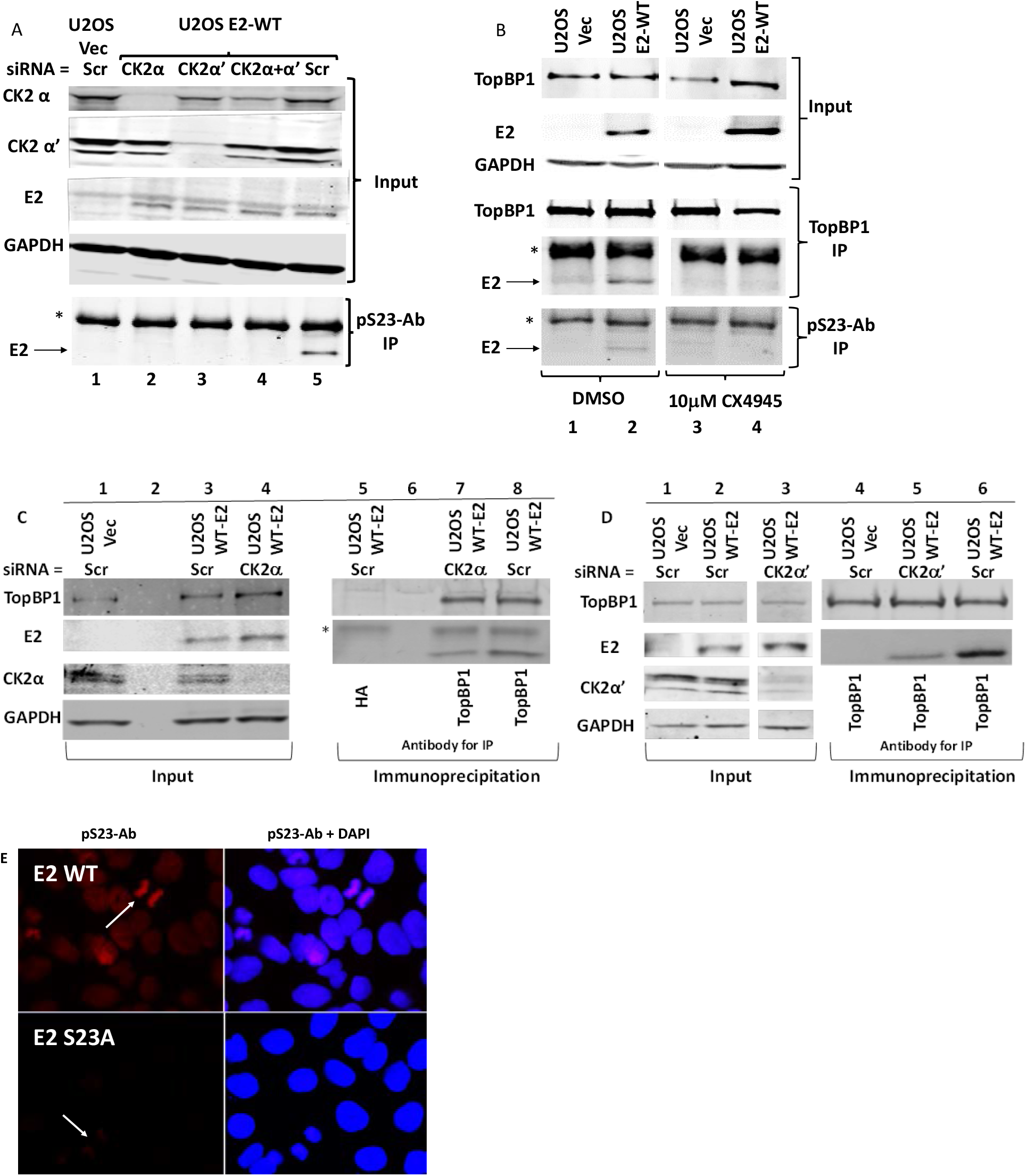
A. siRNA knockdown of CK2α and/or CK2α’. Scr control siRNA was used in lanes 1 and 5. The top panels demonstrate the input proteins that were used in the immunoprecipitation (IP) with pS23-Ab (an antibody raised against an E2 peptide containing a phosphorylated serine 23). Please note the CK2α blot is independent of the other inputs but is run with the same protein extracts. The IP was blotted for E2 which is clearly detected in the Scr control, lane 5, but not in the CK2 knockdown cells, lanes 2-4). B. The indicated cells were treated with DMSO (lanes 1 and 2) or the CK2 inhibitor CX4945 (lanes 3 and 4) and western blotting details the levels of TopBP1 and E2 in the treated cells (top panels). Immunoprecipitation (IP) with TopBP1 demonstrates a pull-down of TopBP1 and E2 (lane 2) that is abrogated by CX4945 (lane 4), middle panels. IP with pS23-Ab pulled-down E2 in control cells (lane 2) that was abolished by CX4945 (lane 4). C. CK2a siRNA knockdown (lane 4) disrupted the E2-TopBP1 interaction as there was a reduced co-IP of E2 in the absence of CK2α (lane 7). D. CK2α’ knockdown (lane 3) disrupted the E2-TopBP1 interaction as there was a reduced co-IP of E2 in the absence of CK2α’ (lane 5). Please note that lanes 1-3 are from the same gel and the same exposure with a lane removed. E. Staining of U2OS E2-WT (top panels) and U2OS E2-S23A (bottom panels) with pS23-Ab. Left hand panels are antibody only, right hand panels are antibody plus DAPI. There was no signal generated with secondary only antibody, and no signal detected in U2OS-Vec control when the primary antibody was included. Figures S2A-B summarizes quantitation for repeat experiments of C and D, respectively. * is an antibody band.

To confirm that E2 is phosphorylated on serine 23 during mitosis, we prepared mitotically enriched U2OS E2-WT and E2-S23A cells. Figure 3E demonstrates a strong signal in the E2-WT mitotic and interphase cells following pS23-Ab staining. With E2-S23A cells, staining with pS23-Ab generates no visible staining in interphase cells, and a very marginal signal in mitotic cells. Figure 5 clearly shows that the E2-S23A protein is detectable by immunofluorescence in these cells with a non-phospho specific E2 antibody, and Figure 1B demonstrates equivalent expression levels of E2-WT and E2-S23A in U2OS cells. Overall, Figures 2 and 3 demonstrate that CK2 phosphorylates E2 on serine 23 to promote interaction with TopBP1.

### CK2 phosphorylates E2 serine 23 in human keratinocytes

We extended our studies into human keratinocytes, the natural target cell type for HPV16 infection. We established N/Tert-1 (human foreskin keratinocytes immortalized by hTERT) cells expressing E2-WT and E2-S23A, as described previously, along with pcDNA Vec control (24). Figure 4A demonstrates expression of E2-WT and E2-S23A in the N/Tert-1 cells (compare lanes 2 and 3, respectively, with lane 1). Immunoprecipitation brought down TopBP1 in all lines (lanes 4-6, top panel), but only E2-WT interacted strongly with TopBP1, demonstrating that the E2-S23A mutation also disrupted the E2-TopBP1 interaction in N/Tert-1 cells (compare lane 6 with lane 5). Figure 4B demonstrates that the pS23-Ab recognizes E2-WT in N/Tert-1 cells; IP with pS23-Ab pulls down E2 (lane 2). The addition of CX4945 (the CK2 inhibitor) to the N/Tert-1 cells abolished E2-WT pull-down with pS23-Ab (compare lane 5 with lane 2), demonstrating that CK2 is responsible for the phosphorylation of E2 serine 23 in N/Tert-1 cells. Abrogation of E2 serine phosphorylation by CX4945 disrupted the E2-TopBP1 interaction (Figure 4C; the inputs from Figure 4B were used). TopBP1 pulled-down both E2-WT and E2-S23D (as it does in U2OS cells, Figure 1), lanes 2 and 3. Treatment with CX4945 disrupted the interaction between TopBP1 and E2-WT but had no effect on the interaction with TopBP1 interaction with E2-S23D. Overall, these results demonstrate that the E2-TopBP1 interaction in N/Tert-1 cells is mediated by CK2 phosphorylation of serine 23 in N/Tert-1 cells.

**Figure 4.**
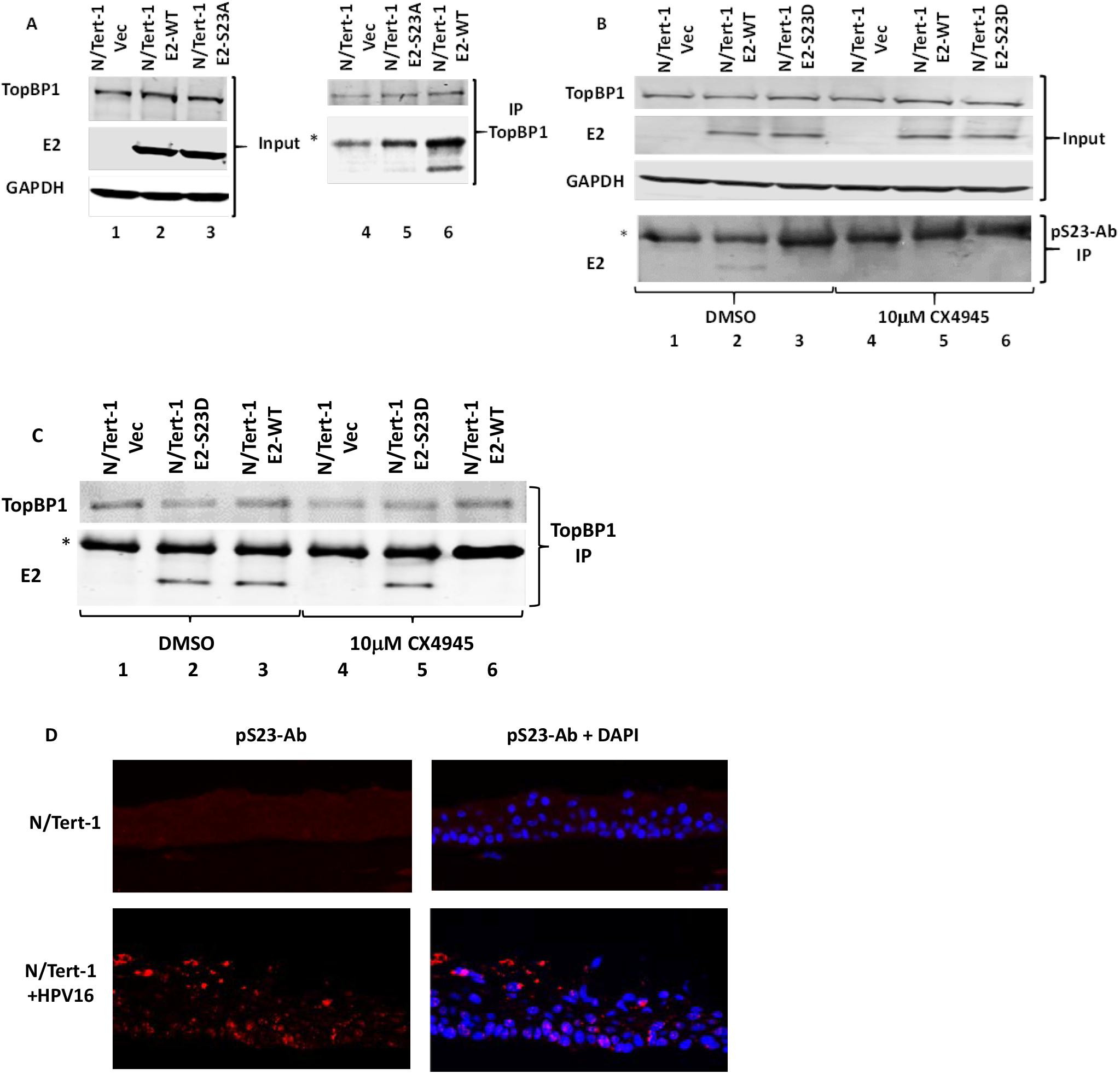
A. Lanes 1-3 is a western blot of extracts from the indicated stable N/Tert-1 cell lines. Lanes 4-6 are western blots of a TopBP1 immunoprecipitation (IP) of the indicated extracts. TopBP1 co-IPs E2-WT but not E2-S23A. B. The extracts in the top panels (Input) were immunoprecipitated with pS23-Ab and E2 is pulled-down by this antibody (bottom panel, lane 2). The CK2 inhibitor CX4945 abrogates this pull-down (lane 5). C. The extracts from B were immunoprecipitated with TopBP1 and both E2-WT and E2-S23D co-IP with TopBP1 (lanes 2 and 3). Treatment with the CX4945 abrogates the interaction between TopBP1 and E2-WT (lane 6), but not E2-S23D (lane 5). D. Organotypic raft cultures of N/Tert-1 (top panels) and N/Tert-1+HPV16 (bottom panels) were stained with pS23-Ab. There is no specific staining in N/Tert-1 cells, but clear staining in N/Tert-1+HPV16. Left panels, pS23-Ab only, right panels, pS23-Ab plus DAPI staining. * is an antibody band.

To investigate whether E2 S23 is phosphorylated during the HPV16 life cycle, we stained N/Tert-1 and N/Tert-1+HPV16 organotypic raft cultures with the pS23-Ab. Our previous work demonstrates that N/Tert-1+HPV16 cells have episomal HPV16 genomes and support late stages of the HPV16 life cycle (24, 81). Figure 4D demonstrates that E2 is phosphorylated on serine 23 in the N/Tert-1+HPV16 cells (bottom panels). The N/Tert-1-Vec cells serve as an isogenic control line, there is no positive signal generated with the pS23-Ab in these cells (top panels). Staining in N/Tert-1+HPV16 is detected throughout the epithelial layer and is clearly nuclear in many of the cells. Please note that the staining outside of nuclei likely relates to nuclear breakdown in the upper layers of the differentiated epithelium, and also potentially reflect ‘smear’ artifacts introduced during micro-sectioning. There is positive staining that is nuclear in many of the cells in N/Tert-1+HPV16.

### The E2 S23A mutation disrupts E2 plasmid retention function

E2 has three clear roles in the viral life cycle; regulation of transcription from the viral and host genomes, replication of the viral genome in association with E1, and segregation of the viral genome into daughter cells where it acts as a bridge between the viral and host genomes during mitosis. The latter function guarantees that the viral genomes are retained in daughter nuclei following mitosis. We measured the transcriptional activation potential of E2-WT and E2-S23A using our ptk6E2-luc system, a plasmid with 6 E2 sites located upstream from the HSV-1 tk promoter driving expression of luciferase (82, 83). Both E2-WT and E2-S23A were able to activate transcription from this reporter with no significant difference in activity (Figure S3A). E2 can repress transcription from the HPV16 long control region (LCR) and E2-WT and E2-S23A were equally able to repress transcription from our pHPV16-LCR-luc reporter ((84) Figure S3B). Using our transient E1-E2 DNA replication assay, we demonstrated that both E2-WT and E2-S23A were able to activate replication with no significant difference between them ((85) Figure S3C).

We then investigated the role of the E2-TopBP1 interaction in plasmid segregation/retention. For this function, E2 interacts with mitotic chromatin. U2OS cells have excellent nuclear architecture also support HPV replication and the maintenance of episomal genomes (86). We synchronized the U2OS cells to enrich for mitotic cells and Figure 5A gives representative images from cells stained with DAPI, E2 and TopBP1. In the absence of E2, TopBP1 was not seen to interact with the mitotic chromatin specifically, instead showing punctate staining, as observed previously (45) (top panels). E2-WT and TopBP1 (middle panels) co-localized on mitotic chromatin, demonstrating that the presence of E2 promotes TopBP1 interaction with the mitotic chromatin. The mutant E2-S23A localized to mitotic chromatin along with TopBP1, but the observed staining was less intense. This pattern of staining was similar in all mitotic cells from the three cell lines. The less intense staining of mitotic chromatin with E2-S23A, when compared with E2-WT, suggests that the interaction between E2 and TopBP1 may elevate E2 protein levels during mitosis. To investigate this, U2OS cells were synchronized by double thymidine blocking and protein harvested from the cells every two hours up until 12 hours following release from blocking. Western blotting was then performed on protein extracts from the cell (Figure 5B). Flow cytometry of the cell lines demonstrating their cell cycle status at different time points are shown in Figure S4. This experiment was repeated and the results for E2 and TopBP1 quantitated (Figure S5). Strikingly, there was a significant increase in E2 and TopBP1 levels 8 hours following release in the E2-WT cells, but an increase of neither E2 nor TopBP1 in E2-S23A or Vec control cells. The 8-hour time point correlates strongly with mitosis (Figure S4). Overall, the results demonstrate that a major role for the E2-TopBP1 interaction is to promote increased levels of both proteins during mitosis. It is also clear that there are other factors that can regulate the interaction of E2 with chromatin as E2-S23A still interacts with mitotic chromatin, and TopBP1 is also recruited to mitotic chromatin with E2-S23A; these points are discussed further below. The increase of E2 and TopBP1 levels during mitosis could contribute to the plasmid segregation/retention function of E2. There were no robust existing assays available for measuring this E2 function, therefore we developed one.

**Figure 5.**
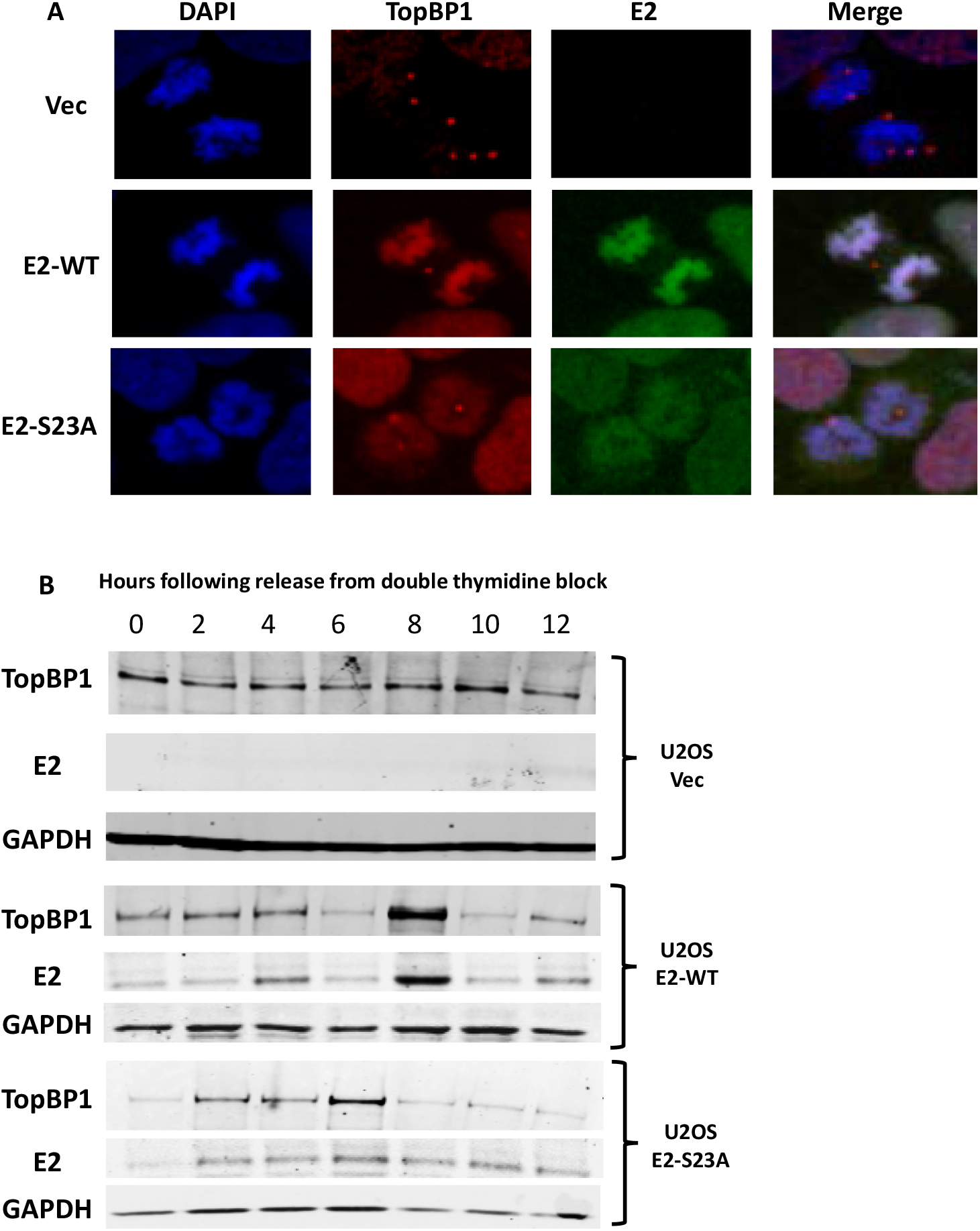
A. U2OS-Vec (top panels), E2-WT (middle panels) and E2-S23A were stained with TopBP1 or E2 as indicated. In the right hand panels a merge of the two antibodies with DAPI staining is shown. TopBP1 does not specifically associate with mitotic chromatin in the Vec cells, but does in the E2-WT and E2-S23A cells. The staining seemed more intense for E2 and TopBP1 with E2-WT than with E2-S23A. These are representative images from experiments where similar phenotypes are repeatedly observed in all three lines B. The indicated U2OS lines were double thymidine blocked to coordinate them in G1. They were then released for the indicated time points, and protein extracts harvested and western blots carried out. At the 8 hour time point there is an enrichment for cells in mitosis in all cell lines (Figure S4). These results were repeated and quantitated and there was a significant increase of E2 and TopBP1 at the 8 hour time point only in U2OS E2-WT cells (Figure S5).

Our stable cell lines allowed us the opportunity to adapt a previously described assay to investigate the plasmid retention function of E2. The assay developed by Silla *et al*. entailed the transient transfection of a plasmid that contained an E2 protein expression cassette along with E2 binding sites driving GFP expression, to allow the measurement of GFP as a readout of E2 plasmid retention function (87). To enhance the quantification of this assay, we removed the E2 expression from the plasmid under retention investigation. This was done by transiently transfecting a plasmid containing E2 binding sites upstream from the tk promoter that drives expression of luciferase, into U2OS cells stably expressing E2 (ptk6E2-luc, Figure 6A summarizes the plasmid retention assay). As a control, cells were transfected with the plasmid pGL3-CONTROL (Promega, pSV40-luc), which contains the SV40 promoter and enhancer driving expression of luciferase activity but has no E2 binding sites present. After 3 days cells were trypsinized and half of each pool were analysed for luciferase activity, while the other half were re-plated. This process was repeated at day 6, whereas at day 9 all of the cells were harvested for luciferase assays. Without selective pressure added to transfected cells, the transfected plasmid DNA is rapidly lost (88). We hypothesized that E2-WT will retain luciferase expression over the 9 days in cells transfected with ptk6E2-luc, while E2-S23A would not. For this assay to work, E2-WT and E2-S23A must be similarly able to activate transcription from ptk6E2-luc, Figure S3A details that this is indeed the case.

**Figure 6.**
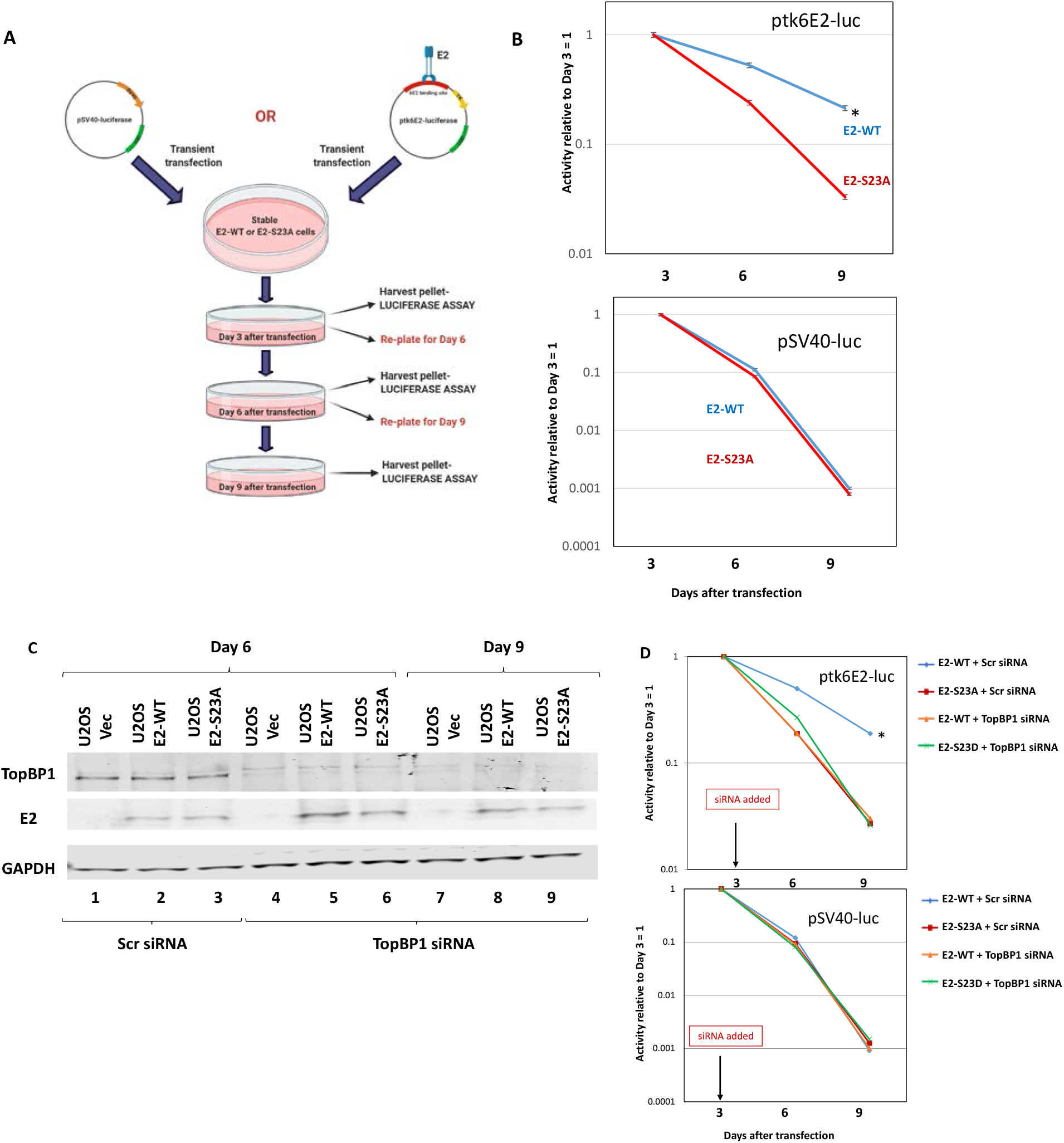
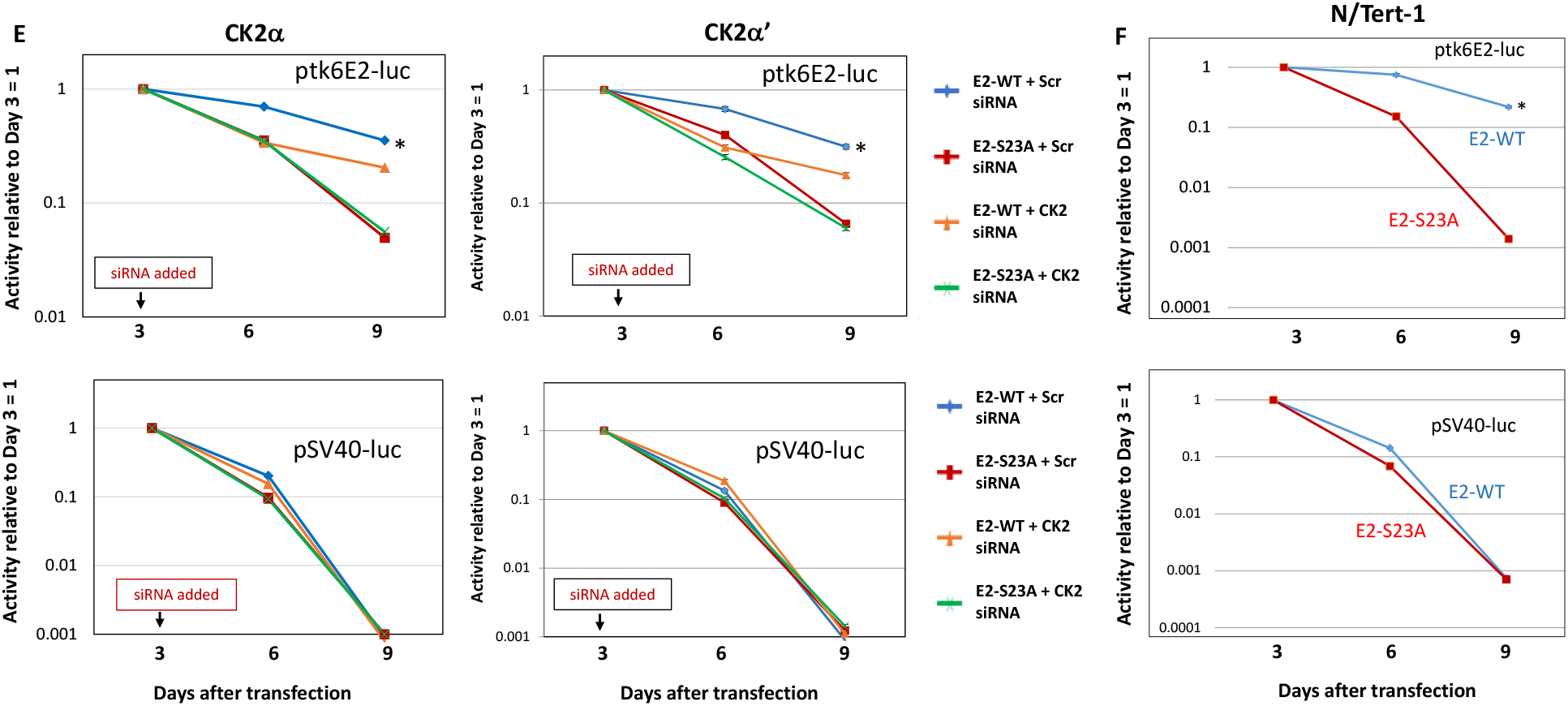
A. A summary of our plasmid retention assay, see text for details. B. In U20S cells, E2-WT retains luciferase expression of ptk6E2-luc while E2-S23A does not (top panel). Neither E2 protein retains expression of pSV40-luc (bottom panel). This is a summary of three independent experiments carried out in duplicate. C. siRNA knockdown of TopBP1 persists until day 9 in our experiment when introduced at day 3. The TopBP1 knockdown (top panel) did not significantly effect the levels of E2 protein throughout the experiment. The siRNA knockdown of TopBP1 did not significantly effect the growth of the cells during the experimental period (Figure S6A). D. The knockdown of TopBP1 with siRNA abolished the ability of E2-WT to retain ptk6E2-luc (top panel), Scr control siRNA had no effect on E2-WT retention (top panel). Neither siRNA treatment changed the expression of pSV40-luc in the experiment (bottom panel). This is a summary of three independent experiments carried out in duplicate. Figure S6B demontrates similar results with alternative TopBP1 siRNAs. E. siRNA knockdown of CK2α or CK2α’ partially disrupted ptk6E2-luc retention by E2-WT (top panels), while having no effect on pSV40-luc activity (bottom panels). This is a summary of three independent experiments carried out in duplicate. Figure S7 demonsrates that the CK2 component expression remains knocked-down throughout the plasmid retention assay. F. In N/Tert-1 cells, E2-WT retains luciferase activity of ptk6E2-luc while S23A does not (top panel). Neither E2 protein retains expression of pSV40-luc (bottom panel). This is a summary of three independent experiments. * indicates a significant difference between the E2-WT and E2-S23A luciferase activity at the day 6 and 9 time points, p-value < 0.05.

Figure 6B demonstrates a significant retention of luciferase activity in E2-WT cells transfected with ptk6E2-luc when compared with transfected E2-S23A cells (note the log_10_ scale). At day 6 E2-WT retained around 80% of the luciferase activity when compared with the day 3 sample, and around 15% at day 9. E2-S23A expressing cells display a hugely reduced luciferase activity retention of around 15% at day 6 and 3% at day 9. The difference in retention between E2-WT and E2-S23A is significant at both day 6 and day 9. With the pSV40-luc control there was no significant difference in luciferase activity between the E2-WT and E2-S23A cells at any time point. As the tk6E2-luc plasmid has minimal activity without the presence of E2, Vec control cells (vector control, no E2 expression) could not be used in this assay.

This assay allows us to measure the retention of nuclear plasmid DNA, as it is based on transcription. A conventional PCR approach would be unable to differentiate between DNA that is being degraded, DNA that is stuck to the cells (or in a cellular compartment other than the nucleus), and DNA that is stuck to the tissue culture substrate. It is possible that E2-S23A fails to retain the plasmid due to the failure to interact with a protein that is not TopBP1. To address this, we took the following approach: Following the day 3 luciferase sampling point, cells were treated with siRNA targeting TopBP1 (siRNA TopBP1-A in materials and methods) and harvested at day 6 and 9 as described. Figure 6C demonstrates effective knockdown of TopBP1 at the day 6 and day 9 time points. The TopBP1 knockdown cells were still growing as determined by a growth curve at the day 6 and 9 time points (Figure S6A). Figure 6D demonstrates that, following siRNA knockdown of TopBP1, the ability of E2-WT to retain luciferase activity is lost. This was reproducible with two additional TopBP1 siRNA targeting sequences (Figure S6B), demonstrating that the defect in E2-S23A during this assay, when compared with E2-WT, is due to the failure to interact with TopBP1. Altogether, the plasmid retention assay showed that E2-WT is able to retain ptk6E2-luc activity in cells via a mechanism that is lost when S23 is altered (E2-S23A), and interaction with TopBP1 abrogated. Figure 6E demonstrates that the knockdown of either CK2α or α’ reduces the ability of E2 to retain luciferase expression of ptk6E2-luc. In both cases there is a sharp reduction in retention of luciferase activity (day 6), but at day 9 there is a partial loss of activity when compared with the E2-S23A mutant activity. This may be because, in the absence of either component, the other begins to compensate to restore active CK2 activity in the cells. CK2α and α’ expression levels remain reduced for the duration of the assay following siRNA treatment (Figure S7). Finally, we demonstrate that our plasmid retention assay functions in N/Tert-1 cells, again demonstrating that E2-WT retains ptk6E2-luc activity while E2-S23A does not. Neither of the E2 variants retain pSV40-luc activity (Figure 6F).

### E2 serine 23 and the HPV16 life cycle

To investigate the role of serine 23 during the HPV16 life cycle, we generated HPV16 genomes containing the E2 S23A and S23D mutations (HPV16-WT, HPV16-S23A, HPV16-S23D). We transfected these genomes into three independent human foreskin keratinocyte (HFK) primary cell cultures to generate immortalized cell lines for life cycle studies. On the first attempt at immortalization, two out of three donors transfected with HPV16-WT and HPV16-S23D generated successful, immortalized cell lines, whereas none of the donors were successfully immortalized by the HPV16-S23A variant (not shown). In the second attempt, we optimized our immortalization procedure by including feeder cells during transfection and selection. Again, we got a striking phenotype with HFK+HPV16-S23A: all three keratinocyte cultures exhibited an attenuated initial immortalization, with slow growing colonies. Figure 7A shows an example for one of the HFK cultures two weeks following selection, crystal violet staining reveals the reduction in colony formation with HPV16-S23A when compared with HPV16-WT and HPV16-S23D. Crystal violet staining following establishment was carried out in duplicate for all three lines and the results are summarized in Figure 7B.

**Figure 7.**
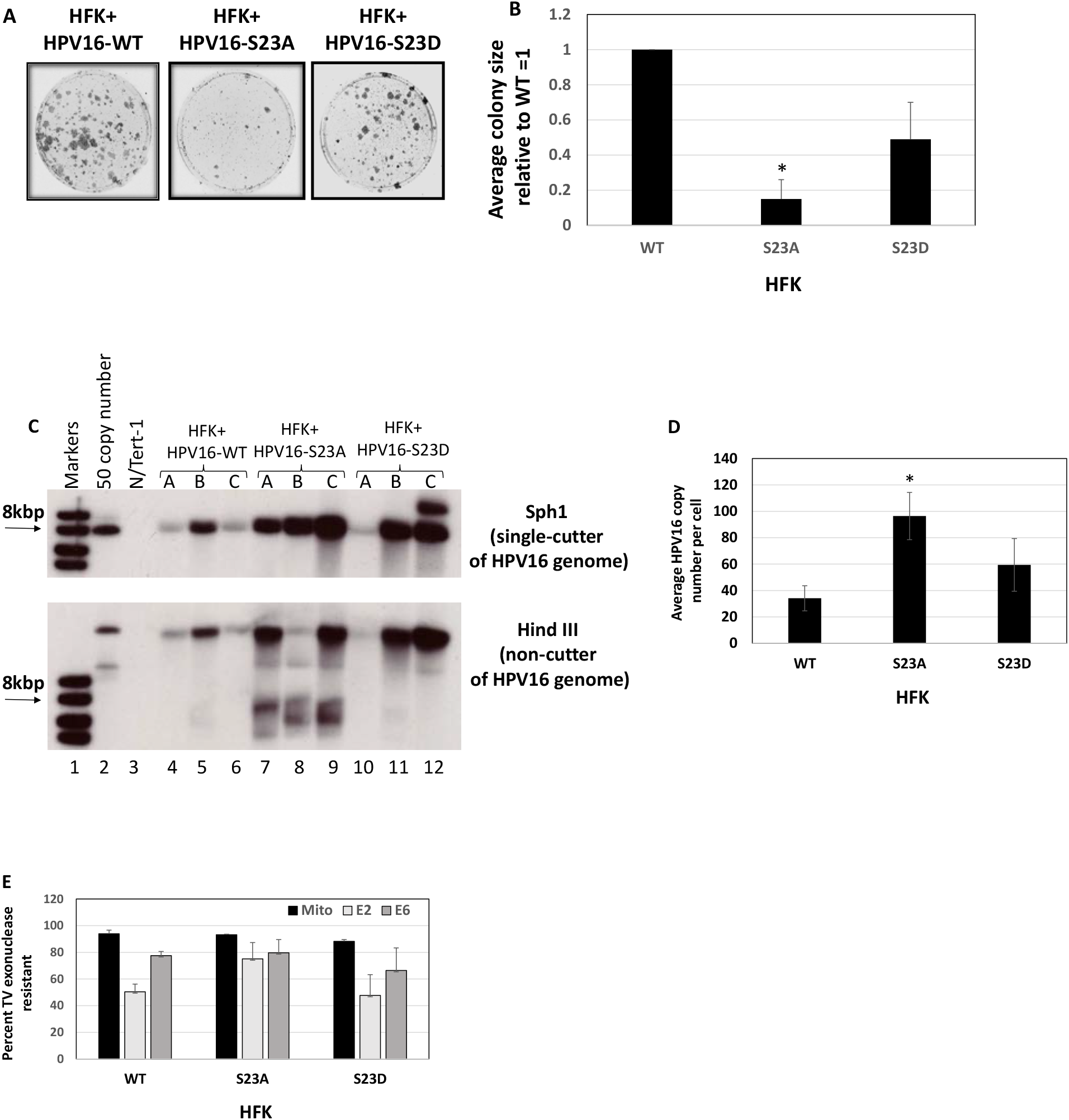
A. Human foreskin keratinocytes (HFK) were transfected with the indicated HPV16 genomes and cell colonies formed 2 weeks after transfection and selection with G418 (pcDNA3 G418 resistant plasmid colonies did not grow out, nor did non-transfected controls). There was a clear reduction in colony size in three independent HFK donors transfected with HPV16-S23A. This was quantitated and the results shown in B. * indicates a significant reduction in colony size for E2-S23A, p-value < 0.05. C. DNA extracted from the indicated cell lines were probed with the HPV16 genome in Southern blots. The top panel demonstrates the presence of 8kbp bands in all samples following digest with the HPV16 genome single cutter Sph1. In the bottom panel results with Hind III, which does not cut the HPV16 genome, are shown. In all cases there is a higher molecular weight band that runs similarly to the control DNA (lane 2). D. The bands in the top panel of C (Sph1 cut) were quantitated and are summarized here. * indicates a significant increase in HPV16-S23A genome copy number, p-value < 0.05. E. To investigate the episomal status of the HPV16 genomes in all cell lines we used the TV exonuclease assay, see text for details. There was no significant difference in the episomal status of the HPV16 genomes between E2-WT, E2-S23A or E2-S23D. Table S1 details the results. Following establishment, the growth rates of the HFK+HPV16-WT, HFK+HPV16-S23A and HFK+HPV16 S23D in monolayer cell culture were similar (Figure S8).

Even though initial immortalization was attenuated, HFK+HPV16-S23A cells eventually grew out successfully. Their growth rate was no different from HFK+HPV16-WT or HFK+HPV16-S23D (Figure S8). To determine the status of the HPV16 genomes in the cells (episomal or integrated) Southern blot analysis was carried out (Figure 7C). In all of the HPV16 lines, *Sph*I (top panel, cuts the HPV16 genome once) generated an 8 kbp signal. Lane 3 contained DNA from N/Tert-1 cells and generated no signal. A band of around 10kbp was observed in HFK+HPV16-S23D-3. A *Hin*dIII (bottom panel, does not cut the HPV16 genome) digest generated a slowly migrating species in all samples, indicative of open circular DNA. Cell lines from all three donors containing HPV16-S23A samples, exhibit significantly faster migrating bands when compared with cells containing WT and S23D genomes (compare lanes 7-9 with the others). A striking feature of the *Sph*I digest is that there is more DNA present in the HFK lines containing the S23A variant when compared with the WT (compare lanes 7-9 with 4-6). The signals generated in the HPV16 lines were quantitated relative to the 50 copy number (lane 2) in the *Sph*I digest. Figure 7D summarizes the quantitation; there is a statistically significant increase in HPV16 genome copy number in the S23A samples compared to WT.

These HFK lines represent pools, therefore there could be increased integration events in some of the lines compared with others. If there were a general level of increased integration, DNA would be integrated in millions of different sites and would not generate detectable signals on Southern blots. We used a recently developed technique that uses exonuclease resistant DNA as a measure of episomal status to determine whether mutation of E2 S23 alters the episomal/integrated status of the HPV16 genomes (89-91). In this assay DNA is treated with TV exonuclease (Exo) which degrades linear DNA, but not circular. We used GAPDH as our linear standard and designated the dCt between samples plus and minus ExoV as 100% degradation. We then took the dCt for a mitochondrial marker (a circular genome) and E2 and E6 and determined the percent of degradation by comparing the dCt difference with that of GAPDH. The data shown are a summary of the three cell lines generated (Figure 7E). The circular mitochondrial DNA is around 90% resistant in all samples. E2 and E6 are between approximately 50 and 80% resistant. Table S1 summarizes the results from these assays. Interestingly, HFK+HPV16-S23D-C is predominantly integrated, and the additional band on the Southern blot (Figure 7D, lane 12) may be related to this. What is clear from this experiment is that there is not a significant difference between HFK+HPV16-WT and HFK+HPV16-S23A/S23D with regards the episomal status of the viral genomes. Therefore, the introduction of these mutations do not promote integration of the viral genome into that of the host.

## Discussion

Here we demonstrate that CK2 phosphorylation of E2 serine 23 results in complex formation with TopBP1 *in vitro* and *in vivo*. The conservation of serine 23 across α-HPV types (Figure 1), and the negative aspartic and glutamic acid residues at −1 and −3 respectively, indicate a potential CK2 target residue (76). *In vivo*, mutation of S23 to alanine disrupts the co-immunoprecipitation of E2 with TopBP1, while an aspartic acid mutation (giving a negative charge mimicking phosphorylation) retains interaction (Figure 1). To demonstrate phosphorylation of S23 *in vivo* we generated a phospho-specific antibody (pS23-Ab) which recognizes E2-WT, including during mitosis, but not E2-S23A (Figure 3). CK2 functions as a tetramer with two β-subunits and two α or α’-subunits, the latter being the enzymatic components of the complex (76). In Figure 3, knockdown of either α component, or partial knockdown of both, abolished detectable levels of E2 S23 phosphorylation in U2OS cells and partially disrupted the interaction between E2 and TopBP1. The reason for the complete loss of phosphorylation, and only a partial loss of interaction, could be due to the failure to detect residual E2 phosphorylation following knockdown of the CK2 components. Unfortunately, pS23-Ab did not work on western blots, it is possible the antibody only recognizes native E2 and not the denatured versions generated for western blotting. Another result supporting the important role for CK2 in the phosphorylation of S23 in U2OS cells is that addition of the CK2 inhibitor CX4945 abolished detectable phosphorylation on this residue (as determined by pS23-Ab immunoprecipitation), and also disrupted the E2-TopBP1 interaction (Figure 3). We extended our studies in U2OS cells to demonstrate that S23 is critical for the E2-TopBP1 interaction in N/Tert-1 cells, and that CX4945 abolishes detectable phosphorylation of E2 on this residue and blocks the E2-TopBP1 interaction in N/Tert-1 cells. We also observed detectable E2 S23 phosphorylation during the HPV16 life cycle in N/Tert-1 cells (Figure 4).

As well as demonstrating that CK2 phosphorylation of S23 mediates the E2-TopBP1 interaction *in vivo*, we also demonstrated that CK2 controls this interaction *in vitro* (Figure 2). E2-WT cannot interact with TopBP1 *in vitro*, while E2-S23D can. Incubation of E2-WT with CK2 promotes the interaction between E2 and TopBP1 recombinant proteins, and this can be reversed by treatment with lambda phosphatase. As E2 has been shown to interact with CK2 components (78), this latter treatment is important as it demonstrates that the enzymatic function of CK2 is required to promote the E2-TopBP1 interaction and that it does not act as a “bridge” to bring the two proteins together. Despite extensive efforts, we were not able to express and purify recombinant E2-S23A to use as a control in these studies. However, the combination of *in vivo* and *in vitro* results demonstrates that CK2 phosphorylation of E2 S23 is crucial for E2-TopBP1 complex formation.

Previous studies have identified several TopBP1 domains that interact with phosphorylated peptides (80). The region around E2 S23 does not correspond to a consensus sequence for interacting with any of these TopBP1 domains and indeed, an E2 pS23 peptide does not interact with any of these domains (Figure S9). This indicates that E2 interacts with an as yet to be identified domain of TopBP1. Future studies will identify this domain, it will be interesting to determine whether E2 has evolved a unique way to interact with TopBP1 that does not disrupt the ability of TopBP1 to interact with host proteins involved in the DNA damage response, a process important for HPV life cycles (69).

Previous studies have implicated CK2 in several aspects of papillomavirus functions. CK2 phosphorylation of bovine papillomavirus 1 (BPV1) E2 on 301 regulates the stability of this protein (92), although this residue is not conserved on HPV16 E2 and both E2-WT and E2-S23A are expressed at relatively equivalent levels in both U2OS and N/Tert-1 cells. CK2 can regulate the DNA binding of BPV and HPV E1 proteins and can control their DNA replication functions (93). CK2 phosphorylates and regulates HPV18 E1 function, and is important in the life cycle of HPV18 and 11 (86, 94). CK2α was the critical component involved in regulating E1, CK2α’ was not involved. CK2 phosphorylation of BRD4 is important for mediating HPV16 E2 transcription and replication function, and we and others have demonstrated that a direct interaction between E2 and BRD4 is required for E2 transcription function (34, 95, 96). As well as regulating E1-E2 functions, CK2 can also regulate the function of E7 proteins. Phosphorylation of a CK2 consensus sequence on E7 is important for E7 degradation of p130 and the promotion of S-phase in differentiated keratinocytes (97), and a HPV18 E7 CK2 target residue is required for maintaining the transformed phenotype of cervical cancer cells (98). Overall, this critical role of CK2 during multiple stages of viral life cycle supports initiatives to investigate the anti-viral and therapeutic effects of CK2 inhibition on HPV infections and disease.

In transient replication and transcription assays the E2-S23A function is similar to E2-WT (Figure S3). A third major function for E2 in the viral life cycle is to mediate viral genome segregation into daughter cells by actively associating with viral and human DNA simultaneously during mitosis (25). E2 has been shown to bind to mitotic chromatin, and previously we demonstrated co-staining of E2 and TopBP1 on mitotic chromatin (37). Therefore, we investigated the interaction of E2-WT and E2-S23A with mitotic chromatin (Figure 5). E2-WT showed robust staining on mitotic chromatin, and in addition it recruited TopBP1 onto the mitotic chromatin. In control cells with no E2, TopBP1 does not “coat” the mitotic chromatin as it does with E2-WT. Therefore, like BPV1 E2 and BRD4, E2 alters TopBP1 interaction with mitotic chromatin (99). For E2-S23A, there was a reproducible reduction in E2 staining on mitotic chromatin although it was located on the mitotic chromatin, and also recruited TopBP1 to the mitotic chromatin. Cell cycle analysis demonstrates that E2-WT levels are increased during mitosis, while E2-S23A levels are not, agreeing with the staining patterns observed. Additionally, E2-WT increases the levels of TopBP1 during mitosis while E2-S23A cannot. Therefore, E2-WT and E2-S23A have distinct phenotypes during mitosis. We are currently investigating whether E2 is stabilized during mitosis. We next investigated whether this difference could contribute to the ability of E2 to retain plasmids in the U2OS cells. Figure 6 demonstrates that E2-WT retains E2 binding site plasmids over an extended period, while E2-S23A could not. A non-E2 binding site plasmid could not be retained by either E2 protein. siRNA knockdown of TopBP1 abrogated the ability of E2-WT to retain the E2 binding site plasmid in this assay. Knockdown of CK2α or α’ also compromised the plasmid retention function of E2-WT, further demonstrating the critical role CK2 plays in mediating the E2-TopBP1 interaction. These results demonstrate that an interaction between E2 and TopBP1 is critical for the plasmid retention function of E2-WT. It is intriguing that E2-S23A retains interaction with mitotic chromatin. Previous work has demonstrated that interaction with mitotic chromatin in itself is not enough to mediate plasmid retention function by E2, two properties seem to be required (87). One property for BPV1 E2 is transcriptional activation (that is mediated by BRD4), and the other is not known. We propose that there is an additional factor that mediates the interaction of E2-S23A with mitotic chromatin and this is under active investigation, BRD4 is clearly a leading candidate we are focusing on.

We introduced S23A and S23D mutations into the HPV16 genome and demonstrated that the E2-S23A genomes are delayed in their ability to immortalize human foreskin keratinocytes (HFK). In addition, the resultant immortalized cell lines that eventually grow out from the HPV16-S23A transfected cells have increased episomal viral genomes when compared with HPV16-WT. Perhaps, at the early stages of establishing immortalization, the viral genome segregation function of E2 is critical to spread the viral genome to daughter cells, and with the S23A mutant genomes this is not occurring. This may delay the growth of the immortalized cells. This process may also explain the increase in viral genome copy number with the S23A mutant as the viral genomes may be mis-segregated into a smaller number of cells, resulting in the ultimate growth of immortalized HFK that have increased viral genomes. It is noticeable that, in the HFK+HPV16-S23A immortalized cell lines, there is a significant increase in faster mobility genomes when compared with HFK+HPV16-WT on Southern blots where the viral genomes have not been cut (Figure 7). During mitosis, TopBP1 is required for decatenation of the host genome to promote the correct segregation of the host chromosomes into daughter nuclei (72, 100). It is possible that the E2-TopBP1 interaction plays a similar role for the circular viral genome during mitosis, ensuring that individual viral genomes reside in the daughter nuclei and are therefore substrates for replication. Therefore, the faster migrating genomes observed in Southern blot analysis of HFK+HPV16-S23A cells could be those that remain catenated. This defect in decatenation could also delay the growth of immortalized HFK. During monolayer cell growth of the HFK lines, it is noticeable that all lines grew equally well (Figure S8). This would suggest that the viral genomes are not being lost in HFK+HPV16-S23A during this period, as loss of viral genomes would result in a reduction in proliferation. This type of growth of the HFK does not represent a “real” stage of the viral life cycle as HPV16 must go through its life cycle in infected human skin. Therefore, during the culture of the HFK in monolayer perhaps the E2-TopBP1 plasmid retention mechanism is not required to maintain viral genome copy numbers. Future studies will focus on more in depth life cycle studies to determine the contribution of the E2-TopBP1 interaction to multiple stages of the viral life cycle.

We have demonstrated that E2 regulates host gene transcription that is relevant to the viral life cycle (24, 101). TopBP1 regulates transcription of E2F and p53 and therefore interaction with E2 could disrupt this process (102, 103). Recently, it has also been demonstrated that TopBP1 regulates host gene transcription during keratinocyte differentiation (104). Finally, while we cannot detect any difference in the replication functions of E2-WT and E2-S23A in our transient replication assays, it is possible that the S23A mutation could compromise replication function of E2 during immortalization. We are actively investigating the functional roles of E2 during the viral life cycle to determine what E2 phenotype of S23A contributes to the aberrant aspects of the viral life cycle already identified.

In summary, we have demonstrated that CK2 phosphorylation of E2 on serine 23 promotes interaction with TopBP1, and that this is important during the viral life cycle. Phenotypically, disrupting this interaction abrogates the plasmid retention function of E2. Future studies will focus on a full understanding of the E2 phenotypes responsible for mediating the life cycle defects observed with the S23 mutants. We also propose that CK2, as it is involved in multiple steps in the HPV16 life cycle, represents a therapeutic target for either blocking or treating HPV16 infections and diseases.

## Materials and methods

### Generation and culture of stable cell lines

Stable cell lines expressing wild type E2 (E2-WT), E2-S23A and E2-S23D, along with pcDNA empty vector plasmid control were established both in U2OS and N/Tert-1 cell lines as previously described (23, 24). Cell culture was also as described in these publications.

### Western blotting

Protein from cell pellets was extracted with 2x pellet volume protein lysis buffer (0.5% Nonidet P-40, 50mM Tris [pH 7.8], and 150mM NaCl) supplemented with protease inhibitor (Roche Molecular Biochemicals) and phosphatase inhibitor cocktail (Sigma). The cells were lyzed for 20 min on ice followed by centrifugation at 18,000 rcf (relative centrifugal force) for 20 min at 4°C. Protein concentration was estimated colorimetrically using a Bio-Rad protein assay. 50 µg of protein with equal volume of 4X Laemmli sample buffer (Bio-Rad) was denatured at 95°C for 5 min. The samples were run on a Novex™ WedgeWell™ 4 to 12% Tris-glycine gel (Invitrogen) and transferred onto a nitrocellulose membrane (Bio-Rad) using the wet-blot method, at 30 V overnight. The membrane was blocked with *Li-Cor* Odyssey® blocking buffer (PBS) diluted 1:1 v/v with PBS and then incubated with specified primary antibody in *Li-Cor* Odyssey® blocking buffer (PBS) diluted 1:1 with PBS. Following this, the membrane was washed with PBS supplemented with 0.1% Tween20 and further probed with the Odyssey secondary antibodies (IRDye® 680RD Goat anti-Rabbit IgG (H + L), 0.1 mg or IRDye® 800CW Goat anti-Mouse IgG (H + L), 0.1 mg) in *Li-Cor* Odyssey® blocking buffer (PBS) diluted 1:1 with PBS at 1:10,000 for 1 h at room temperature. After washing with PBS-tween, the membrane was imaged using the Odyssey^®^ CLx Imaging System and ImageJ was used for quantification. Primary antibodies used for western blotting studies are as follows: HPV16 E2 (TVG 261) or monoclonal B9 1:500 (Abcam ab17185 for TVG261, (105) for monoclonal B9), TopBP1 1:1000 (Bethyl; catalog no. A300-111A), GAPDH 1:10,000 (Santa Cruz; catalog no. sc-47724), Casein kinase IIα (1AD9) 1:500 (Santa Cruz; catalog no. sc-12738), CKII alpha’ Antibody 1:1000 (Bethyl; catalog no. A300-199A).

### Immunoprecipitation

Primary antibody of interest or a HA-tag antibody (used as a negative control) was incubated in 250 µg of cell lysate (prepared as described above), made up to a total volume of 500 µl with lysis buffer (0.5% Nonidet P-40, 50mM Tris [pH 7.8], and 150mM NaCl), supplemented with protease inhibitor (Roche Molecular Biochemicals) and phosphatase inhibitor cocktail (Sigma) and rotated at 4°C overnight. The following day, 40 µl of pre-washed protein A beads per sample (Sigma, equilibrated to lysis buffer as mentioned in the manufacturer’s protocol) was added to the lysate:antibody mixture and rotated for another 4 hours at 4°C. The samples were gently washed with 500 µl lysis buffer by centrifugation at 1,000 rcf for 2-3 min. This wash was repeated 4 times. The bead pellet was resuspended in 4X Laemmli sample buffer (Bio-Rad), heat denatured and centrifuged at 1,000 rcf for 2-3 min. Proteins were separated using an SDS-PAGE system and transferred onto a nitrocellulose membrane before probing for the presence of E2 or TopBP1, as per the western blotting protocol.

### Immunofluorescence and cell synchronization

U2OS cells expressing stable E2-WT, E2-S23A and pcDNA empty vector plasmid control were plated on acid-washed, *poly-L-lysine* coated coverslips, in a 6-well plate at a density of 2 × 10^5^ cells / well (5 ml DMEM + 10% FBS media). After 24 h, the cells were treated with 2 mM thymidine diluted in the DMEM-FBS for 16 h. This was then washed 2 times with PBS and recovered in supplemented DMEM. After 8 h, to block the cells at G1/S phase, a second dose of 2 mM thymidine was added and incubated for 17 h. The cells were then washed twice with PBS and recovered as before for 3 h. The cells were next treated with nocodazole (100 ng/ml) for 5 h and released for 2 h to enrich for mitotic cells. Following this, the cells were washed twice with PBS, fixed and stained as described in (34). The primary antibodies used are as follows: HPV16 E2 (TVG 261) 1:500 (Abcam; ab17185), HPV16 E2 B9 monoclonal antibody, 1:500 (105), TopBP1 1:1000 (Bethyl; catalog no. A300-111A), pS23-Ab 1:10,000 (Custom generated by GenScript; peptide sequence-CKILTHYENDS^P^TDLR). The cells were washed and incubated with secondary antibodies Alexa fluor 488 goat anti-mouse (Thermo fisher; catalog no. A-11001) and Alexa fluor 594 goat anti-rabbit (Thermo fisher; catalog no. A-11037) diluted at 1: 1000. The wash step was repeated, and the coverslips were mounted on a glass slide using Vectashield mounting medium containing 4′,6-diamidino-2-phenylindole (DAPI). Images were captured with a Zeiss LSM700 laser scanning confocal microscope and analyzed using Zen LE software.

### Cell synchronization

U2OS cells expressing stable E2-WT, E2-S23A and pcDNA empty vector plasmid control were plated at 3 × 10^5^ density onto 100-mm plates in DMEM + 10% FBS media. The cells were treated with 2 mM thymidine diluted in the supplemented DMEM media for 16 h. The cells were then washed 2 times with PBS and recovered in supplemented DMEM media. After 8 h, to block the cells at G1/S phase, a second dose of 2 mM thymidine was added and incubated for 17 h. The cells were then washed twice with PBS and recovered as before at the following time points: 0 h and 2 h (G1/S phase), 4 h and 6 h (S phase), 8 h (M1 phase), 10 h (M2 phase), and 12 h (the next G1 phase). The cell lysate was prepared using the harvested cells at different time points and immunoblotting was carried out as described.

### Plasmid segregation assay

This assay is based upon the well-established fact that transfected DNA is lost from cells if there is no selective pressure put on them to retain the plasmid DNA; transfected DNA is quickly lost from the cells after 48-72 hours (88). Two luciferase reporter plasmids were used for our novel assay; one containing the SV40 promoter and enhancer (pGL3 Control, Promega, described as pSV40-luc in this manuscript) which has no E2 DNA binding sites, the other with the HSV1 tk promoter driving expression of luciferase with 6-E2 target sites upstream (82, 83). The pSV40-luc and the ptk6E2-luc were transiently transfected, separately into either E2-WT or E2-S23A cells. 3 days post-transfection, the cells were trypsinized and half re-plated and half harvested for a luciferase assay system (Promega). This luciferase activity was considered as the baseline activity. At day 6, the same process was repeated, half of the cells was harvested for luciferase assay, half re-plated. At day 9, cells were harvested solely for luciferase activity assay. Bio-Rad protein estimation assay was used for protein concentration estimation, to standardize for cell number. Relative fluorescence units were measured using the BioTek Synergy H1 hybrid reader. The activities shown are expressed relative to the respective protein concentrations of the samples. The assays shown are representative of three independent experiments carried out in triplicates.

### Small interfering RNA (siRNA) and segregation assay

U2OS parental cells were plated on a 100-mm plates. The next day, cells were transfected with 10 µM of following siRNA. 10 µM of MISSION^®^ siRNA Universal Negative Control (Sigma-Aldrich; catalog no. SIC001) was used as a “non-targeting” control in our experiments. Lipofectamine™ RNAiMAX transfection (Invitrogen; catalog no. 13778-100) protocol was used in the siRNA knockdown. 48 h post transfection, the cells were harvested, and knockdown confirmed by immunoblotting for the protein of interest. Segregation assays were performed as described before, after treating the cells with the siRNA of interest on day 3 of the protocol.

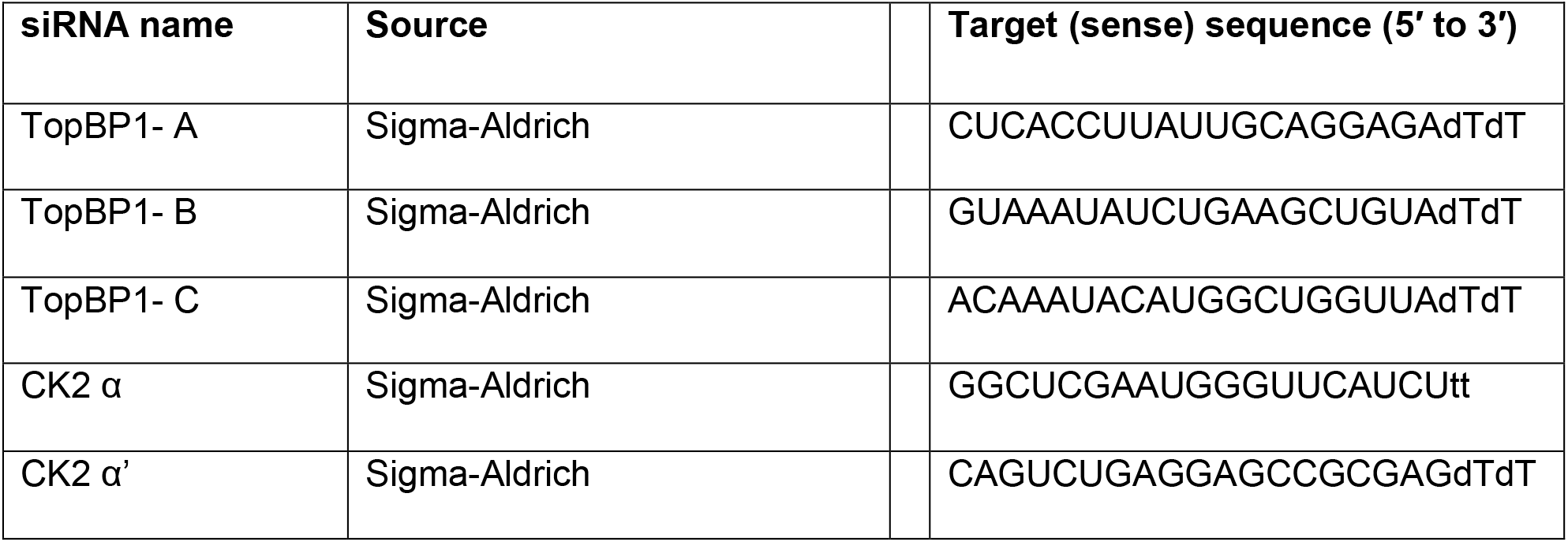

### Production of recombinant protein

Amino-acids 1-200 of E2-WT and E2-S23A were produced as fused protein with His-tag, and TopBP1 was produced as a fused protein with a GST-tag (GST TopBP1 (aa 32-1522) His from Addgene; plasmid # 20375). The protein expression was carried out by picking a single colony of BL21(DE3) Competent *E. coli* (NEB Inc.; catalog no. C2527) and growing it in LB media supplemented with 100 *μ*g/ml of selective antibiotics (kanamycin for His-tagged E2-WT and E2-S23D; ampicillin for GST-tagged TopBP1), grown overnight at 37°C and shaken at a low speed. This starter culture was then diluted 1:100 in fresh LB media with kanamycin. The culture was shaken at 37°C until the optimal density of 0.6-0.8 at OD600n was achieved. Following this, IPTG at final concentration of 1 mM was added to the culture, for induction of protein expression, shaking at 16°C overnight. His-tagged proteins were purified on Ni-NTA agarose (Qiagen; catalog no. 30761) and GST-tagged TopBP1 protein was purified on Glutathione Sepharose™ 4B (GE healthcare; catalog no. 17-0756), according to the batch purification method described in the manufacturer’s manual, followed by size-exclusion chromatography. The purity of the recombinant protein was confirmed by SDS–PAGE analysis.

### *In vitro* GST pull-down assays

Purified recombinant His-tagged E2-WT and E2-S23D protein and GST-tagged TopBP1 were used for the *In vitro* pull-down assays. GST-tagged NEDD4 E3 ligase was used as our GST-control. GST-TopBP1 and GST-control were kept stable at 0.65 pmol and 11 pmol of His-E2-WT and His-E2-S23D were used for the experiment. Glutathione Sepharose™ 4B (GE healthcare; catalog no. 17-0756), equilibrated to the GST lysis buffer (50 mM Tris-HCl, pH 8.0, 100 mM NaCl, 0.5 mM EDTA, 0.5 mM EGTA and 0.5% NP-40, 1 mM DTT plus protease inhibitors) was added and each tube placed at 4ºC for 1 h with continual end-to-end rotation. The protein bound GST beads were washed 3 times in the GST lysis buffer by centrifugation at 1,000 rcf for 3 minutes and resuspended in 4X Laemmli sample buffer (Bio-Rad), heat denatured and centrifuged at 1,000 rcf for 3 minutes. The supernatant was gel electrophoresed using an SDS-PAGE system which was later transferred onto a nitrocellulose membrane using wet-blot transfer method. The membrane was probed for the presence of E2 or TopBP1, as described above.

### *In vitro* kinase assay with or without lambda phosphatase

Immunoprecipitated GST beads were prepared as mentioned above in the GST-pull down section. After 1 h, the beads were incubated with 1 µl CK2 enzyme and 1X CK2 reaction buffer (NEB Inc; catalog no. P6010S) supplemented with 200 µM ATP and 30 mM MgCl2 and rotated for 1 h at 30ºC. The beads were then incubated in presence or absence of lambda phosphatase (Santa Cruz; catalog no. sc-200312A) as described in the manufacturer’s protocol. Following this, the beads were washed and analyzed by immunoblotting.

### CK2 inhibitor treatment

U2OS and N/Tert-1 cells were plated at a density of 2 × 10^5^ in a 100-mm plate. Next day, the cells were treated with 10 µM CK2 inhibitor, CX-4945 (Silmitasertib) from APExBIO (catalog no. A8330) or 10 µM DMSO for 48 h. The cells were then harvested and processed for immunoprecipitation with pS23Ab or TopBP1 as described.

### Immortalization of human foreskin keratinocytes (HFK)

HPV16 mutant genomes (S23A and S23D) were generated by Genscript. The HPV16 (WT, S23A, S23D) were removed from their parental plasmid using Sph1, and the viral genomes isolated and then re-circularized using T4 ligase (NEB) and transfected into early passage HFK from three donor backgrounds (Lifeline technology), alongside a G418 resistance plasmid, pcDNA. Cells underwent selection in 200 ug/mL G418 (Sigma-Aldrich) for 14 days and were cultured on a layer of J2 3T3 fibroblast feeders (NIH), which had been pre-treated with 8 μg/ml mitomycin C (Roche). Throughout the immortalization process, HFK were cultured in Dermalife-K complete media (Lifeline Technology). In Figure 5A, transfected cells were stained with crystal violet 14 days following transfection and selection prior to passaging.

### Southern blotting

Total cellular DNA was extracted by proteinase K-sodium dodecyl sulfate digestion followed by a phenol-chloroform extraction method. 5 ug of total cellular DNA was digested with either SphI (to linearize the HPV16 genome) or HindIII (which does not cut the HPV16 genome). All digests included DpnI to ensure that all input DNA was digested and not represented as replicating viral DNA. All restriction enzymes were purchased from NEB and utilized as per manufacturer’s instructions. Digested DNA was separated by electrophoresis of a 0.8% agarose gel, transferred to a nitrocellulose membrane, and probed with radiolabeled (32-P) HPV16 genome. This was then visualized by exposure to film for 1 to 24 hours. Images were captured from an overnight-exposed phosphor screen by GE Typhoon 9410 and quantified using ImageJ.

### Exonuclease V assay

To examine whether viral genomes were maintained as episomes, we carried out an exonuclease V assay, as described by Bienkowska-Haba *et al*. 2018, which determines the resistance of HPV16 genomes to exonuclease V. 20 ng genomic DNA was either treated with exonuclease V (RecBCD, NEB), in a total volume of 30 ul, or left untreated for 1 hour at 37°C followed by heat inactivation at 95°C for 10 minutes. 2 ng of digested/undigested DNA was then quantified by real time PCR using a 7500 FAST Applied Biosystems thermocycler with SYBR Green PCR Master Mix (Applied Biosystems) and 100 nM of each primer in a 20 μl reaction. Nuclease free water was used in place of the template for a negative control. The following cycling conditions were used: 50°C for 2 minutes, 95°C for 10 minutes, 40 cycles at 95°C for 15 seconds, and a dissociation stage of 95°C for 15 seconds, 60°C for 1 minute, 95°C for 15 seconds, and 60°C for 15 seconds. Separate PCR reactions were performed to amplify HPV16 E6 F: 5’-TTGCTTTTCGGGATTTATGC-3’ R: 5’-CAGGACACAGTGGCTTTTGA-3’, HPV16 E2 F: 5’-TGGAAGTGCAGTTTGATGGA-3’ R: 5’-CCGCATGAACTTCCCATACT-3’, human mitochondrial DNA F: 5’-CAGGAGTAGGAGAGAGGGAGGTAAG-3’ R: 5’-TACCCATCATAATCGGAGGCTTTGG-3’, and human GAPDH DNA F: 5’-GGAGCGAGATCCCTCCAAAAT-3’ R: 5’-GGCTGTTGTCATACTTCTCATGG-3’

## Acknowledgements

This work was supported by VCU Philips Institute for Oral Health Research and the National Cancer Institute Designated Massey Cancer Center grant P30 CA016059 (IMM), Cancer Research UK C6992/A12695 (IMM and BOS), Cancer Research UK C302/A14532 (MD, AWO and LHP) and Cancer Research UK C302/A24386 (MD, AWO and LHP). We thank Dr. Chris Li, VCU Philips Institute for Oral Health Research, for the kind gift of the GST-NEDD4 control plasmid.

